# Contrasting Biological Pathways Underlie Agronomic Traits in Two *Coffea canephora* Breeding Populations

**DOI:** 10.1101/2025.10.31.685932

**Authors:** Ezekiel Ahn, Sunchung Park, Jishnu Bhatt, Seunghyun Lim, Lyndel W. Meinhardt

## Abstract

This study reveals that distinct breeding populations of *Coffea canephora* can achieve agronomic success through fundamentally different biological strategies. To uncover this, we performed a comparative genomic analysis of two populations (‘Premature’ and ‘Intermediate’), integrating single-SNP association, machine learning (Bootstrap Forest), and Gene Ontology (GO) pathway analysis. The genetic architecture of the Premature population was linked to specialized metabolic pathways, including lipid modification and processes within the organelle lumen, a finding supported by the identification of a putative *caffeine synthase 3* gene. In contrast, traits in the Intermediate population were governed by variation in core cellular machinery, with significant enrichment for pathways related to actin cytoskeleton regulation and salicylic acid signaling. This discovery provides a new biological context for important candidate genes involved in disease resistance (e.g., *RPP13*-like, NB-ARC, *CERK1*). These findings demonstrate that population-specific biological routes underpin agronomic performance, providing a powerful foundation for designing more targeted breeding programs in coffee. We release population-specific, ranked SNP lists and GO gene sets as a reusable resource to enable meta-analyses and benchmarking in coffee and perennial crops.

## 1 Introduction

With a long history with the human race, coffee is one of the world’s most valuable agricultural commodities, deeply involved with the economies and livelihoods of millions globally [1]. Global coffee production is dominated by two species: *Coffea arabica* (arabica), renowned for its superior cup quality, and *Coffea canephora* (robusta). While arabica commands a premium, *C. canephora* is a cornerstone of the coffee industry, contributing approximately 40% of global production due to its higher yield [2], inherent disease resistance, and adaptability to a wider range of growing conditions, including warmer temperatures that are increasingly impacting traditional arabica regions [3]. Originating from sub-Saharan Africa, *C. canephora* has a high productivity potential, known to reach 2 to 4 t/ha/year under optimal management strategies, surpassing the 1.5 to 3 t/ha/year typical of *C. arabica* [3]. In many regions, such as Colombia, robusta coffee represents an economic opportunity for small-scale producers. However, like many perennial crops, coffee breeding faces significant challenges, including long breeding cycles, genetically complex traits, and the need for extensive, multi-year field evaluations. These challenges are further complicated by devastating diseases like coffee leaf rust (caused by *Hemileia vastatrix*), which can inflict yield losses of up to 47% in *C. canephora* if left unmanaged [4].

The genetic improvement of coffee through conventional breeding is a notoriously "slow and time-consuming" process [5]. This inefficiency stems from the long juvenile period inherent to perennial crops, which extends the breeding cycle time, a key limiting factor in achieving genetic gain [6,7]. Furthermore, accurately selecting for complex traits like yield and disease resistance, which are heavily influenced by environmental factors, requires expensive, multi-year, and multi-location field trials [8]. These logistical and financial burdens significantly limit the efficiency of traditional phenotypic selection [9]. To accelerate genetic progress, modern genomic tools were developed; however, early methods like Marker-Assisted Selection (MAS) proved less effective for complex traits controlled by many genes with small effects [10]. This has created an urgent need to apply more comprehensive approaches, like Genome-Wide Association Studies (GWAS), to effectively dissect the genetic architecture of these traits in coffee.

The integration of genome-wide markers to predict breeding values has emerged as a powerful strategy for accelerating genetic gains in plant breeding [9]. For perennial species like *C. canephora*, Genomic Selection (GS) is particularly promising, as it has the potential to drastically shorten the long breeding cycle. Indeed, several recent studies have demonstrated the utility of GS in *C. canephora*, confirming its role as a "useful and promising tool" for genetic improvement [10, 11]. However, the routine implementation of GS in coffee breeding programs still faces challenges, including the costs and logistics associated with high-throughput genotyping [13]. While progress has been made in dissecting the genetic basis of key traits in this species, including those in the specific populations analyzed here, these studies often employ different statistical models. A critical need, therefore, exists to compare findings across these diverse analytical frameworks to build a more robust and validated understanding of the genetic architecture of key agronomic traits.

This is especially true for traits related to quality and disease resistance. For instance, the genetic regulation of carotenoid biosynthesis, which influences fruit color and quality, is a key area where candidate genes can be targeted [14]. Likewise, while the genetics of resistance to coffee leaf rust (*Hemileia vastatrix*) have been studied, its complexity warrants further investigation, as genomic studies on disease resistance in coffee are still described as "lacking" [15]. Given that different statistical and machine learning models may identify different candidate genes, a comparative approach is essential to pinpointing the most reliable targets for marker-assisted selection and future functional validation.

This study builds upon the rich genomic datasets and foundational analyses previously conducted in these *C. canephora* populations, which have successfully demonstrated the utility of genomic prediction [16], stability analysis [17], and polygenic association models [18]. While these efforts provide valuable insights, the application of alternative analytical frameworks can offer a more comprehensive understanding of the genetic architecture and help validate key genomic regions. While these studies provided a critical foundation, they primarily focused on the accuracy of predictive models. The underlying biological pathways that drive the distinct genetic architectures of these populations, however, have not been comparatively analyzed and remain a key area for exploration.

Specifically, the foundational work by Ferrão et al. [16] on this dataset focused on comparing the accuracy of various whole-genome statistical models for the purpose of genomic prediction. A subsequent study by Adunola et al. [17] extended this by investigating genotype-by-environment interactions and the predictability of trait stability metrics. While these efforts expertly demonstrated that genomic selection is a viable tool for coffee breeding, the fundamental question of why the genetic control of agronomic traits might differ between these two distinct breeding populations remains unanswered. Our study directly addresses this knowledge gap by shifting the analytical focus from statistical prediction to biological interpretation.

To achieve this, we re-analyze the public data for three key agronomic traits, coffee bean production, leaf rust incidence, and green bean yield, to provide a multi-faceted view of their genetic control. We first identified candidate loci using a conventional single-SNP (single nucleotide polymorphism) regression model to establish a broad set of associations. We then employed a machine learning model, Bootstrap Forest, to provide a complementary perspective by ranking the explanatory contribution of each SNP, allowing us to capture importance from complex, non-linear relationships [19]. By synthesizing the results from these analytical frameworks with a new Gene Ontology (GO) analysis, this work uncovers fundamentally different biological strategies (e.g., specialized metabolism vs. core cellular machinery) that underpin agronomic performance in these populations. This provides a more nuanced understanding of their specific genetic architectures and offers targeted, biologically-informed directions for future functional studies and coffee breeding.

## 2 Materials and Methods

### 2.1 Experimental Populations and Data

As previously mentioned, the data, including both phenotypic & genotypic sets, is sourced from *C. canephora* dataset [16]. Two recurrent selection populations, designated "Intermediate" and "Premature," were developed by the *Instituto Capixaba de Pesquisa*, *Assistência Técnica e Extensão Rural* (*Incaper*), ES State, Brazil. These names refer to their distinct fruit maturation groups; the Premature population ripened and was harvested approximately one month earlier than the Intermediate population [16]. The intermediate population was derived from crosses of 16 progenitors, and the premature population was selected from 9 progenitors for high production of coffee beans and similar stages of fruit maturity. The populations were established in two locations: Marilândia Experimental Farm (FEM; 19°24’ S, 40°31’ W, 70 m altitude) and Sooretama Experimental Farm (FES; 15°47’ S, 43°18’ W, 40 m altitude). Phenotypic data were collected over four consecutive harvest-production years (2008-2011) for three traits: production of coffee beans (mature coffee fruit in the "cherries" stage, in 60-kg bags per hectare); natural infection of coffee leaf rust, caused by *H. vastatrix* (1-9 scale, based on visual sporulation intensity); and yield of green beans post-harvest (ripened beans, in g, after processing). Genotyping was performed using Genotyping-by-Sequencing (GBS), resulting in 45,748 SNPs for the Intermediate population and 59,332 SNPs for the Premature population after quality control. Further details on population development, experimental design, and data collection can be found in Ferrão *et al.* (2019) [16].

### 2.2 Data preprocessing

Phenotypic and genotypic data were sourced from the publicly available dataset established by Ferrão et al. [16]. To ensure direct comparability with previous foundational studies on these populations, we used the final, pre-processed data without modification. The phenotypic values represent adjusted means that account for block and harvest effects as determined by a mixed model. These adjusted means were used directly for all subsequent analyses without further normalization or transformation. The genotypic data had already undergone rigorous quality control, which included the removal of triallelic SNPs, markers with a minor allele frequency (MAF) below 1%, and SNPs with call rates under 70%. SNP genotypes were coded as -1, 0, or 1 to indicate the count of the reference allele and were called using a Bayesian approach that incorporated parental information. All data was imported and managed using JMP Pro 17 (SAS Institute Inc., Cary, NC) [20].

### 2.3 Single-SNP association analysis

We performed a response screening analysis in JMP Pro 17, using the platform’s default settings, to identify individual SNPs associated with each trait. For each population (Premature and Intermediate) and trait combination, the adjusted phenotypic value was used as the response variable, and all SNPs were included as predictor variables. A series of individual linear regressions was performed for each SNP, and a *p*-value for the association was calculated. This approach tests the additive effect of each SNP individually, assuming a linear relationship between the number of reference alleles and the trait value. To control for the high probability of false positives inherent in testing tens of thousands of SNPs, we applied a False Discovery Rate (FDR) correction to the *p*-values. We chose a stringent FDR-adjusted significance threshold of 0.01 to identify a robust set of high-confidence SNP-trait associations.

### 2.4 Machine learning for SNP importance analysis

#### 2.4.1 Model rationale and implementation

To complement the single-SNP linear regressions, which test each marker individually, we employed a machine learning approach to identify important loci by considering all SNPs simultaneously. A preliminary model screening was conducted using JMP Pro 17 to compare the performance of several available algorithms for explaining phenotypic variation (data not shown). The Bootstrap Forest algorithm was selected for all subsequent analyses as it consistently provided the best explanatory performance across the different traits. The Bootstrap Forest algorithm, a robust ensemble method well-suited for high-dimensional genomic data, can capture complex, non-linear, and interactive effects that may be missed by single-marker models [19,21]. Critically, our objective for using this model was not for developing a predictive tool, but for variable importance ranking, a method to identify the SNPs with the greatest explanatory power. Given that the goal was explanatory and not predictive, potential model overfitting for performance on a new dataset was not the primary concern. Instead, the model was used to generate a comprehensive and complementary set of candidate genes for our primary objective: the downstream biological pathway analysis. The analysis was conducted in JMP Pro 17 for each population and trait combination, using all available SNPs as input features.

#### 2.4.2 Model parameters and variable importance

To ensure reproducibility and transparency, the Bootstrap Forest models were built using the following fixed parameters: Number of Trees = 100; Bootstrap Sample Rate = 1; Minimum Splits Per Tree = 10; Maximum Splits Per Tree = 2,000; Minimum Size Split = 5; and a fixed random seed for reproducibility. The Number of Terms Sampled Per Split was adjusted based on the number of SNPs in each population (30,333 for Premature; 23,086 for Intermediate), a high mtry value chosen to prioritize locus discovery over predictive accuracy. Variable importance for each SNP was then quantified using the "Portion" statistic, which represents the relative contribution of each SNP to the model. The top five SNPs with the highest "Portion" values for each trait and population were identified for further analysis.

### 2.5 Candidate gene identification and gene ontology enrichment analysis

To build a comprehensive list of candidate genes for pathway analysis, we leveraged the significant loci identified from our two complementary analytical frameworks: the statistically significant SNPs from the single-SNP association analysis and the top-ranked SNPs from the Bootstrap Forest importance analysis. For each significant SNP identified by the single-SNP association analysis (FDR-adjusted *p*-value < 0.01) and for the top five most important SNPs from the Bootstrap Forest analysis, we located the nearest annotated gene. This was performed using the *C. canephora* genome browser (https://coffee-genome-hub.southgreen.fr/coffea_canephora) [22]. For each identified gene, we recorded its gene ID, putative function, and the distance (in base pairs) from the associated SNP.

To investigate the broader biological pathways underlying the genetic architecture of these traits, we conducted a GO enrichment analysis. For this, a list of candidate genes was generated from the loci of the most important SNPs identified by the Bootstrap Forest models. To capture a robust biological signal for each trait-population combination while excluding SNPs with negligible explanatory power, we selected the genes associated with the top 100 most important SNPs for the enrichment analysis. The enrichment analysis was performed using ShinyGO v0.88 (http://bioinformatics.sdstate.edu/go/) [23] against the *Coffea canephora* gene database (AUK_PRJEB4211_v1), which contains 49,390 background genes. We performed separate analyses for the ’Premature’ and ’Intermediate’ populations, as well as for combined population gene lists, across various trait combinations. The analysis was performed using default settings, with a FDR cutoff of 0.05 for statistical significance and a pathway size filter to include GO terms containing between 2 and 5,000 genes. The complete results of this enrichment analysis for all trait and population combinations are available in Supplementary Data 1.

## 3 Results

### 3.1 Single-SNP association analysis of agronomic traits

To identify SNPs associated with key agronomic traits in *C. canephora*, we performed a response screening analysis examining three traits (coffee bean production, leaf rust incidence, and green bean yield) in two populations (Premature and Intermediate). We applied an FDR-adjusted *p*-value threshold of 0.01 to identify significant SNP-trait associations.

The single-SNP analysis revealed a stark contrast in the genetic architecture between the two populations (Fig. 1). No significant SNP associations were found for the production of coffee beans or the yield of green beans in the Intermediate population (FDR > 0.01).

**Fig 1.**
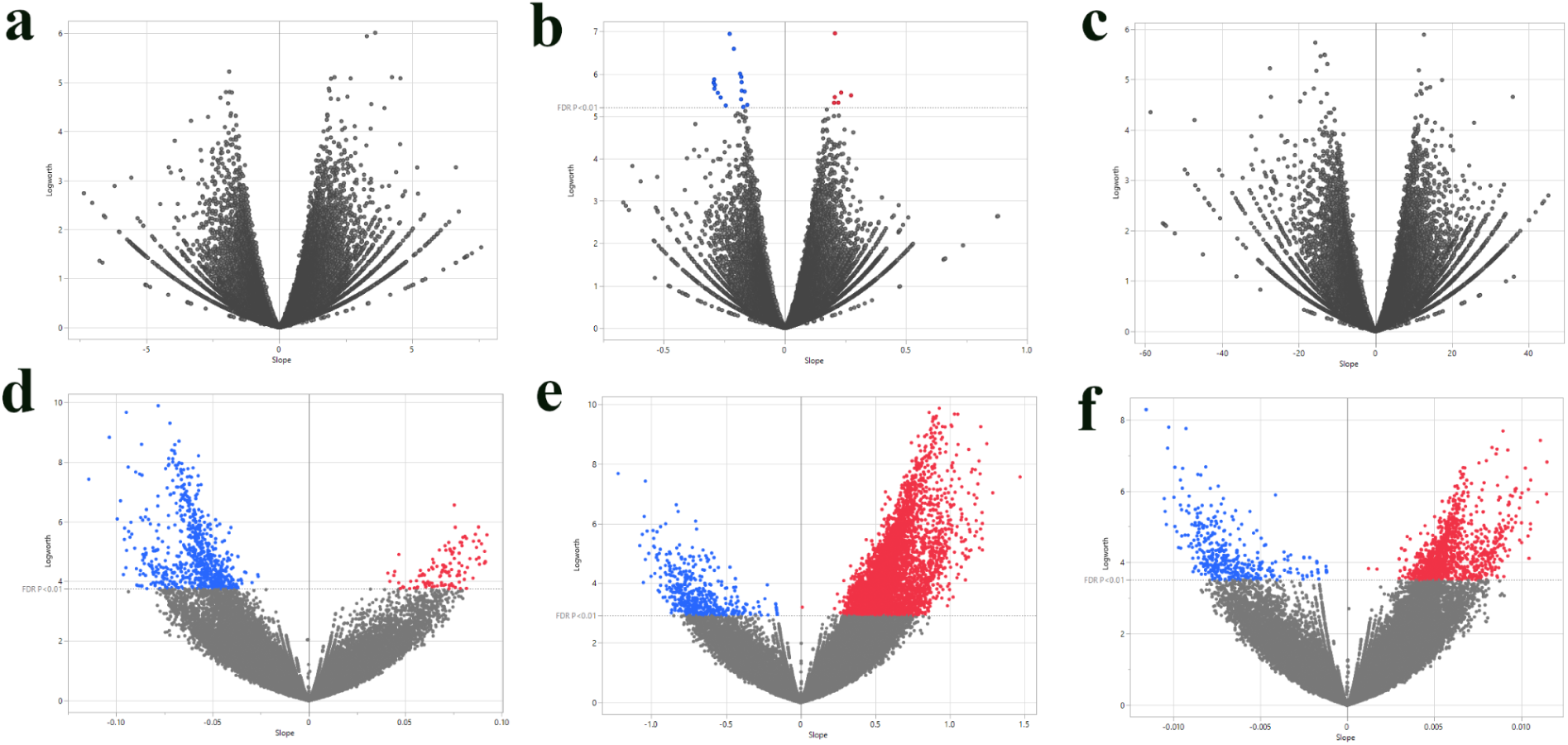
Volcano plots of single-SNP association analysis for three agronomic traits in two *C. canephora* populations. Each point represents a single SNP. The x-axis shows the estimated slope (effect size) from a linear regression of the phenotype on the SNP genotype, and the y-axis shows the negative base-10 logarithm of the *p*-value from the association test. SNPs are colored based on the direction of their effect and statistical significance after applying an FDR correction to account for multiple testing. Red points indicate SNPs with a positive effect and FDR-adjusted *p*-value < 0.01; blue points indicate SNPs with a negative effect and FDR-adjusted *p*-value < 0.01; gray points do not meet the significance threshold. The plots highlight a substantially larger number of significant associations in the Premature population compared to the Intermediate population. (a) Production of coffee beans, Intermediate. (b) Leaf rust incidence, Intermediate. (c) Yield of green beans, Intermediate. (d) Production of coffee beans, Premature. (e) Leaf rust incidence, Premature. (f) Yield of green beans, Premature.

However, for leaf rust incidence in this population, a total of 23 significant SNPs were identified (17 with negative effects and 6 with positive effects) (Fig. 1b). In sharp contrast, the Premature population showed thousands of significant associations across all traits.

Specifically, we identified 1,020 significant SNPs for the production of coffee beans (Fig. 1d), 7,100 SNPs for leaf rust incidence (Fig. 1e), and 1,850 SNPs for the yield of green beans (Fig. 1f).

To visualize the genomic distribution of significant associations, we generated Manhattan plots showing the R-squared values for each significant SNP across the 11 chromosomes of *C. canephora* (Fig. 2). For the production of coffee beans in the Premature population, a prominent peak was observed on chromosome 6 (Fig. 2a). Significant SNPs were distributed across multiple chromosomes for leaf rust incidence in the Premature population, with particularly strong peaks on chromosomes 5 and 7 (Fig. 2b). Significant SNPs were found on chromosomes 2, 5, 9, and 11 for the yield of green beans in the Premature population (Fig. 2c). Leaf rust incidence in the Intermediate population exhibited significant associations on chromosomes 1, 2, 5, and 10 (Fig. 2d).

**Fig. 2.**
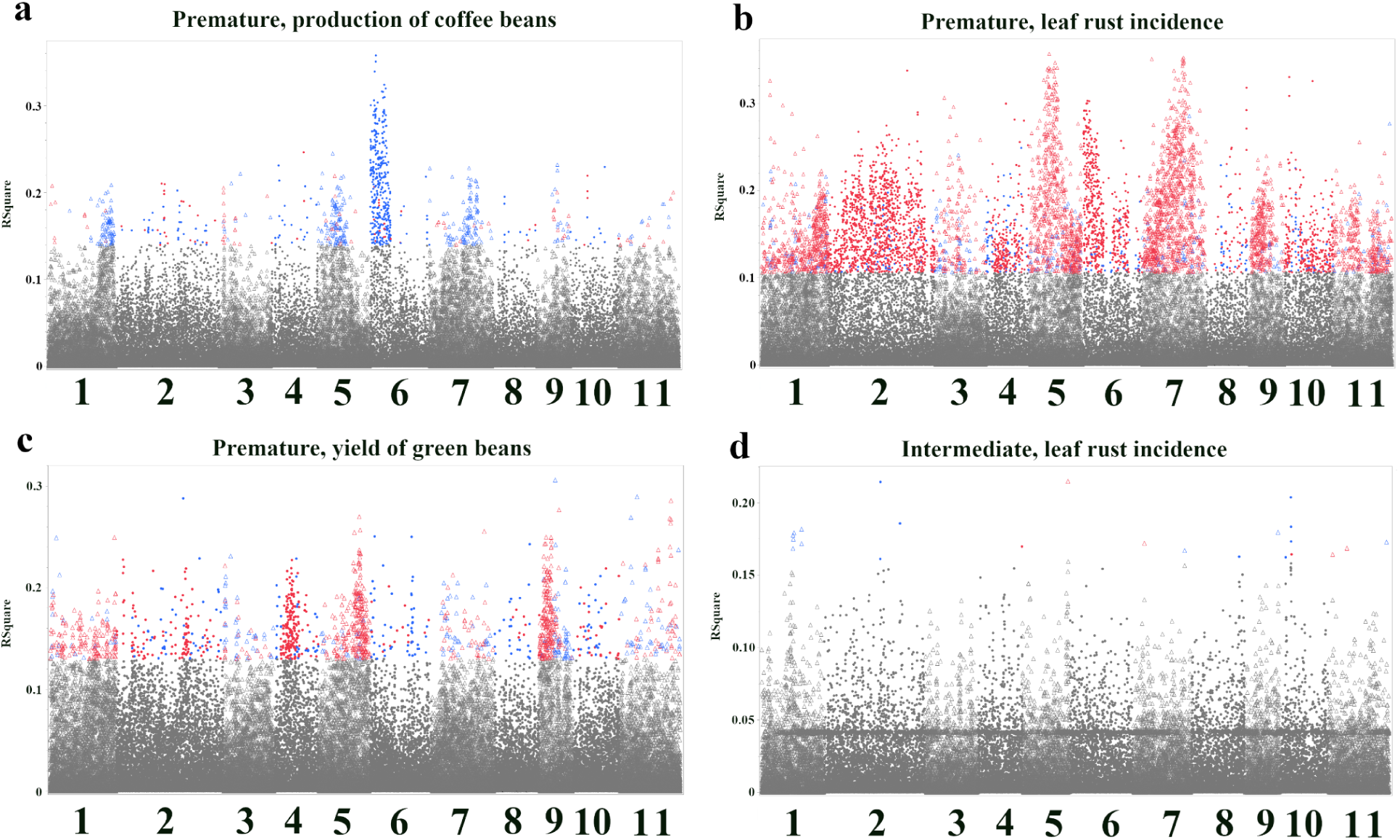
Manhattan plots of single-SNP association analysis, showing only SNPs that reached genome-wide significance (FDR < 0.01). Each point represents a single SNP. The x-axis shows the chromosomal location (1 through 11). The y-axis shows the R-squared value, representing the proportion of phenotypic variance explained by each individual SNP. SNPs are colored based on the direction of their effect: red indicates a positive effect, and blue indicates a negative effect. Each panel reveals the genomic regions most strongly associated with the respective trait. (a) Production of coffee beans, Premature. (b) Leaf rust incidence, Premature. (c) Yield of green beans, Premature. (d) Leaf rust incidence, Intermediate. Plots for two trait-population combinations are not shown due to a lack of significant associations.

Table 1 presents the candidate genes closest to the significant SNPs identified for each population-trait combination. Several of these genes have plausible connections to the traits under investigation. Among the significant SNPs for leaf rust incidence, several genes with known roles in plant defense were identified. In the Premature population, these included a putative disease resistance RPP13-like protein (*Cc04t14270.1*), an NB-ARC domain-containing protein (*Cc03t09860.1*), a peroxidase (*Cc02t30380.1*), and a chitin elicitor receptor kinase 1 (CERK1; *Cc05t03340.1*) (Table 1). RPP13-like and NB-ARC domain-containing proteins are often involved in pathogen recognition and downstream defense signaling. Peroxidases strengthen cell walls and produce reactive oxygen species during defense responses. CERK1 is a key receptor for chitin, a major component of fungal cell walls, triggering immune responses upon pathogen detection. In the Intermediate population, significant SNPs for leaf rust incidence were located near genes encoding a C2H2-type domain-containing protein (*Cc10t04730.1*), a WRKY domain-containing protein (*Cc10t04810.1*), a putative late blight resistance protein homolog R1B-16 (*Cc01t08110.1*), and a RING-type domain-containing protein (*Cc11t03510.1*) (Table 1). WRKY transcription factors are known to regulate plant defense responses [24,25], and RING-type proteins often function as E3 ubiquitin ligases, potentially involved in regulating defense signaling [26].

**Table 1.**
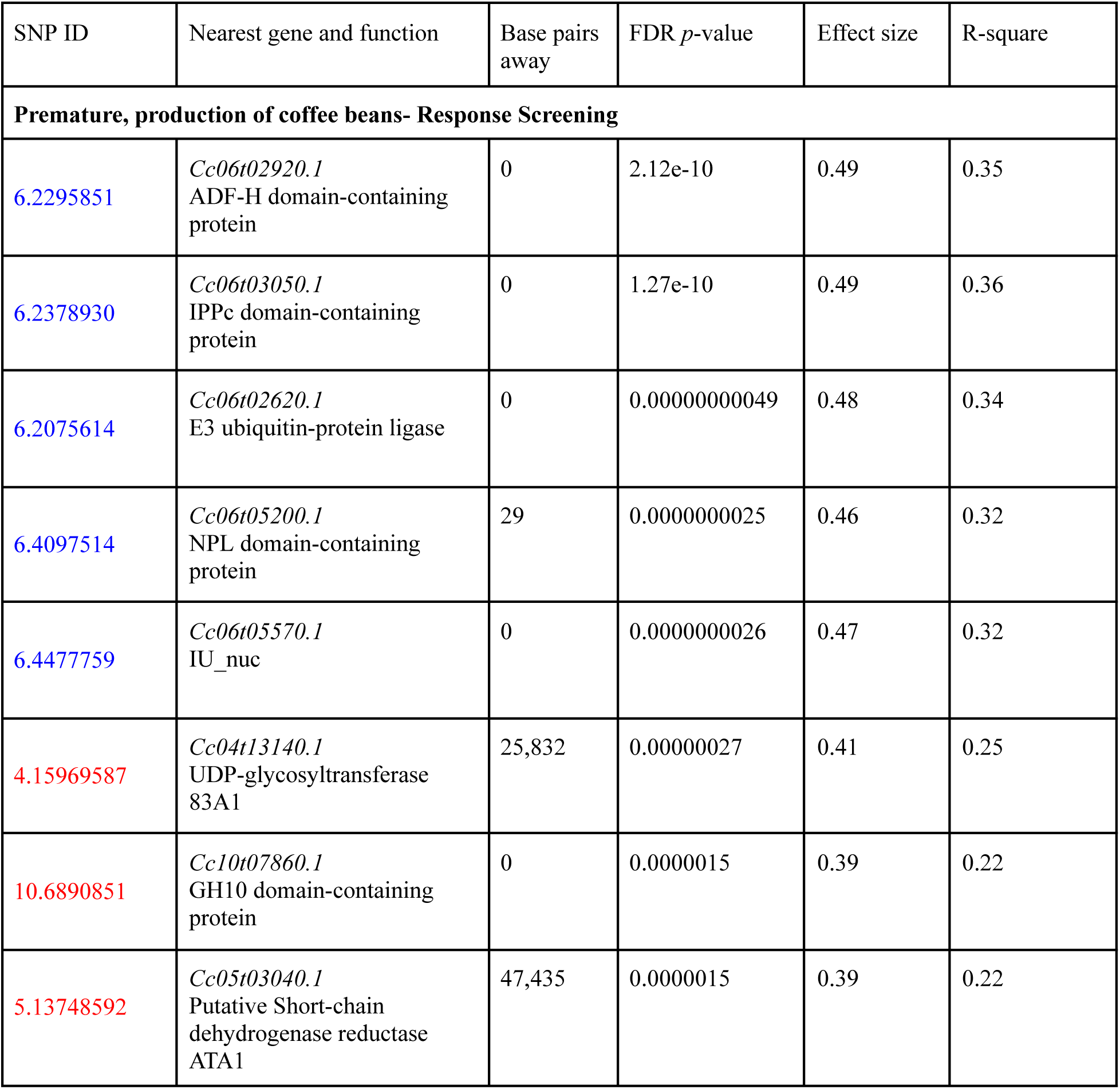
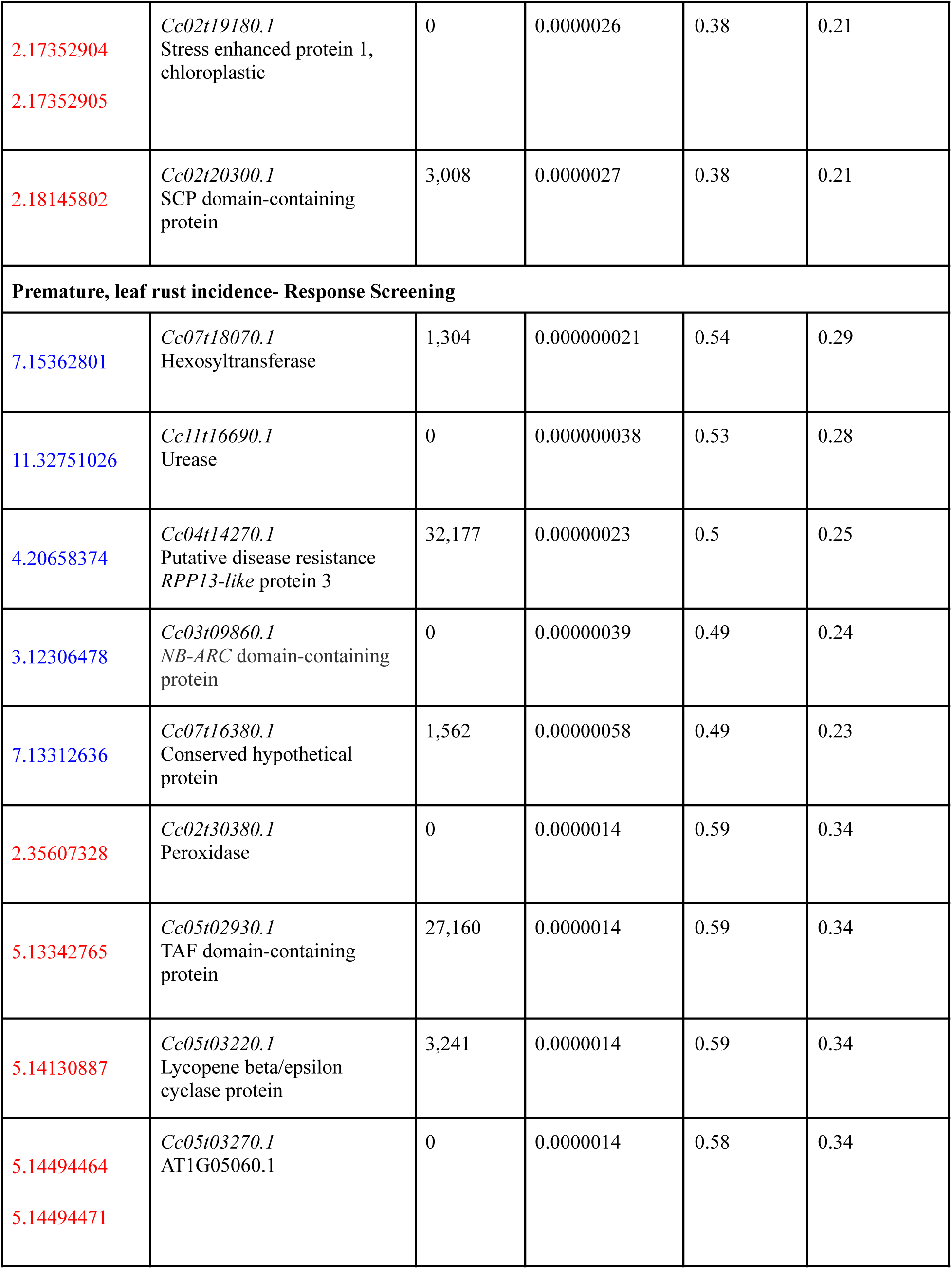
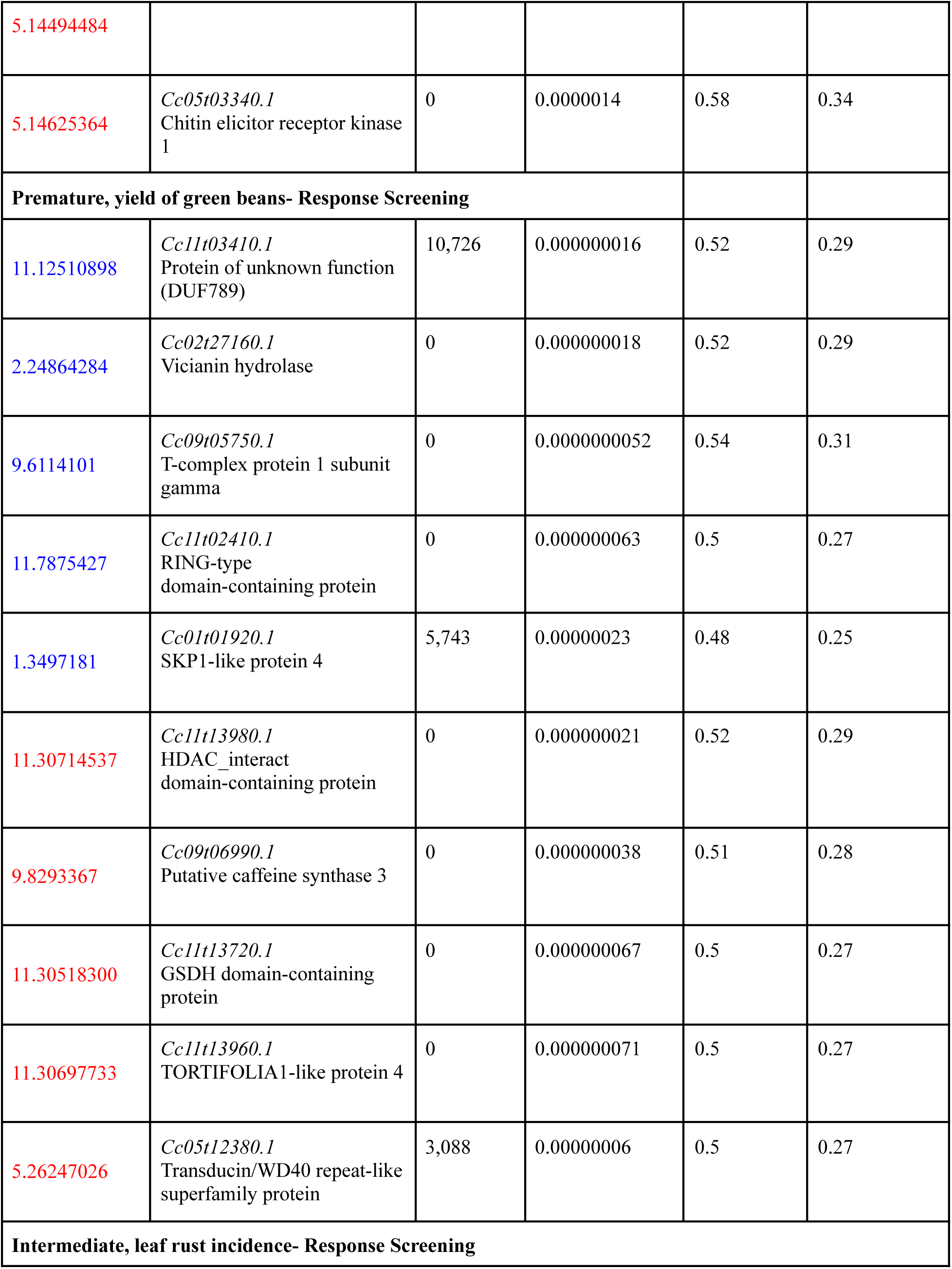
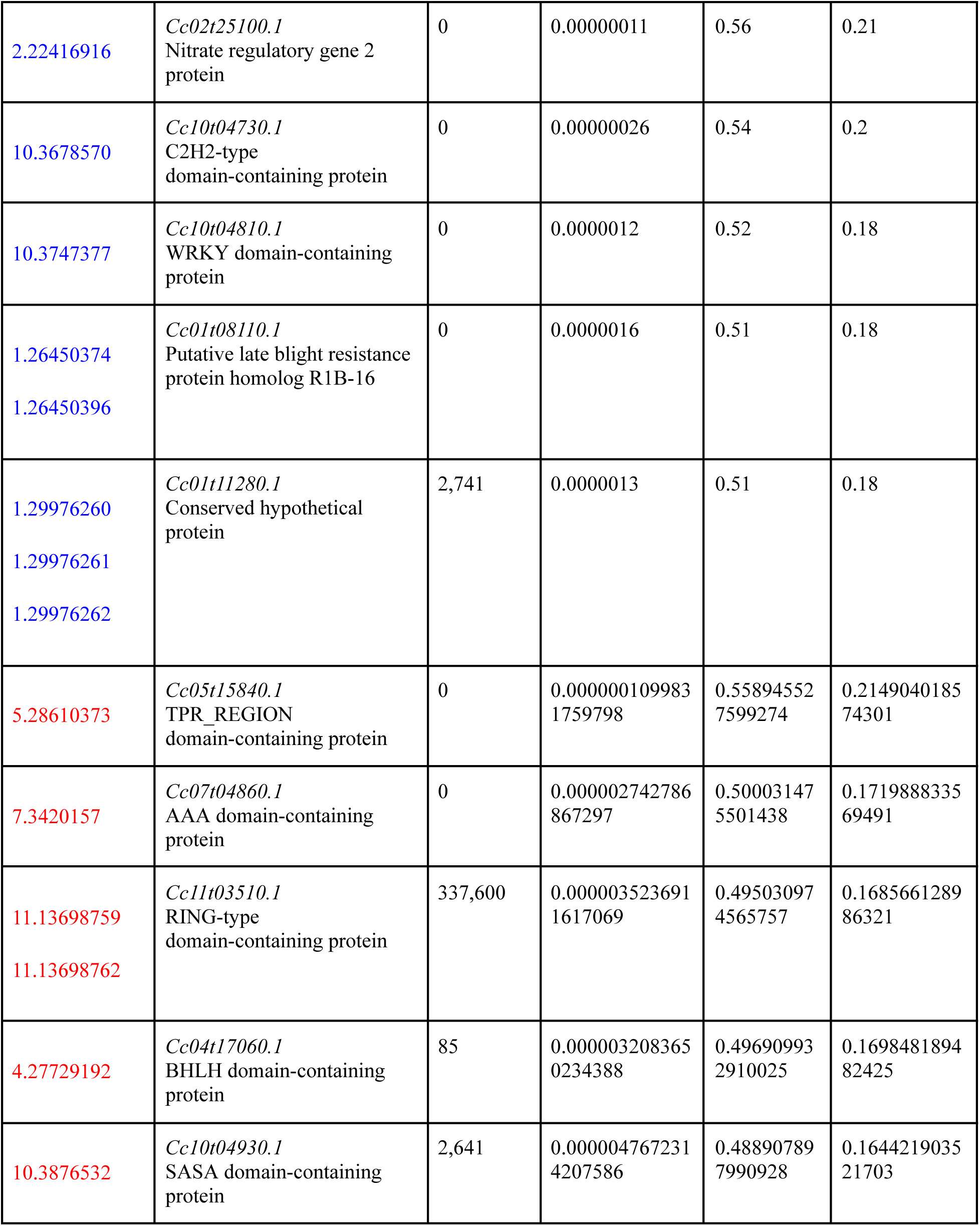
Candidate genes associated with SNPs showing significant associations with agronomic traits in two *C. canephora* populations (Premature and Intermediate). SNPs were identified as significant based on a response screening analysis in JMP Pro 17, with an FDR-adjusted *p*-value threshold of 0.01. For each significant SNP, the table lists: the SNP identifier (SNP ID) (blue= negative and red= positive association); the nearest gene and its putative function (based on the annotation of the *C. canephora* reference genome); the distance in base pairs between the SNP and the start of the nearest gene (0 indicates the SNP is within the gene); the FDR-adjusted *p*-value; the estimated effect size (slope from a linear regression of the phenotype on the SNP genotype); and the R-squared value (proportion of phenotypic variance explained by the SNP). Genes are listed separately for each population and trait combination. Only traits with at least one significant SNP are included.

| SNP ID | Nearest gene and function | Base pairs away | FDR *p*-value | Effect size | R-square |
| --- | --- | --- | --- | --- | --- |
| **Premature, production of coffee beans- Response Screening** |  |  |  |  |  |
| 6.2295851 | *Cc06t02920.1*<br>ADF-H domain-containing protein | 0 | 2.12e-10 | 0.49 | 0.35 |
| 6.2378930 | *Cc06t03050.1*<br>IPPC domain-containing protein | 0 | 1.27e-10 | 0.49 | 0.36 |
| 6.2075614 | *Cc06t02620.1*<br>E3 ubiquitin-protein ligase | 0 | 0.00000000049 | 0.48 | 0.34 |
| 6.4097514 | *Cc06t05200.1*<br>NPL domain-containing protein | 29 | 0.0000000025 | 0.46 | 0.32 |
| 6.4477759 | *Cc06t05570.1*<br>IU_nuc | 0 | 0.0000000026 | 0.47 | 0.32 |
| 4.15969587 | *Cc04t13140.1*<br>UDP-glycosyltransferase 83A1 | 25,832 | 0.00000027 | 0.41 | 0.25 |
| 10.6890851 | *Cc10t07860.1*<br>GH10 domain-containing protein | 0 | 0.0000015 | 0.39 | 0.22 |
| 5.13748592 | *Cc05t03040.1*<br>Putative Short-chain dehydrogenase reductase ATA1 | 47,435 | 0.0000015 | 0.39 | 0.22 |
| 2.17352904<br>2.17352905 | *Cc02t19180.1*<br>Stress enhanced protein 1,<br>chloroplastic | 0 | 0.0000026 | 0.38 | 0.21 |
| 2.18145802 | *Cc02t20300.1*<br>SCP domain-containing<br>protein | 3,008 | 0.0000027 | 0.38 | 0.21 |
| **Premature, leaf rust incidence- Response Screening** |  |  |  |  |  |
| 7.15362801 | *Cc07t18070.1*<br>Hexosyltransferase | 1,304 | 0.000000021 | 0.54 | 0.29 |
| 11.32751026 | *Cc11t16690.1*<br>Urease | 0 | 0.000000038 | 0.53 | 0.28 |
| 4.20658374 | *Cc04t14270.1*<br>Putative disease resistance<br>*RPP13-like* protein 3 | 32,177 | 0.00000023 | 0.5 | 0.25 |
| 3.12306478 | *Cc03t09860.1*<br>*NB-ARC* domain-containing<br>protein | 0 | 0.00000039 | 0.49 | 0.24 |
| 7.13312636 | *Cc07t16380.1*<br>Conserved hypothetical<br>protein | 1,562 | 0.00000058 | 0.49 | 0.23 |
| 2.35607328 | *Cc02t30380.1*<br>Peroxidase | 0 | 0.0000014 | 0.59 | 0.34 |
| 5.13342765 | *Cc05t02930.1*<br>TAF domain-containing<br>protein | 27,160 | 0.0000014 | 0.59 | 0.34 |
| 5.14130887 | *Cc05t03220.1*<br>Lycopene beta/epsilon<br>cyclase protein | 3,241 | 0.0000014 | 0.59 | 0.34 |
| 5.14494464<br>5.14494471 | *Cc05t03270.1*<br>AT1G05060.1 | 0 | 0.0000014 | 0.58 | 0.34 |
| 5.14494484 |  |  |  |  |  |
| 5.14625364 | *Cc05t03340.1*<br>Chitin elicitor receptor kinase 1 | 0 | 0.0000014 | 0.58 | 0.34 |
| **Premature, yield of green beans- Response Screening** |  |  |  |  |  |
| 11.12510898 | *Cc11t03410.1*<br>Protein of unknown function (DUF789) | 10,726 | 0.000000016 | 0.52 | 0.29 |
| 2.24864284 | *Cc02t27160.1*<br>Vicianin hydrolase | 0 | 0.000000018 | 0.52 | 0.29 |
| 9.6114101 | *Cc09t05750.1*<br>T-complex protein 1 subunit gamma | 0 | 0.0000000052 | 0.54 | 0.31 |
| 11.7875427 | *Cc11t02410.1*<br>RING-type domain-containing protein | 0 | 0.000000063 | 0.5 | 0.27 |
| 1.3497181 | *Cc01t01920.1*<br>SKP1-like protein 4 | 5,743 | 0.000000023 | 0.48 | 0.25 |
| 11.30714537 | *Cc11t13980.1*<br>HDAC_interact domain-containing protein | 0 | 0.000000021 | 0.52 | 0.29 |
| 9.8293367 | *Cc09t06990.1*<br>Putative caffeine synthase 3 | 0 | 0.000000038 | 0.51 | 0.28 |
| 11.30518300 | *Cc11t13720.1*<br>GSDH domain-containing protein | 0 | 0.000000067 | 0.5 | 0.27 |
| 11.30697733 | *Cc11t13960.1*<br>TORTIFOLIA1-like protein 4 | 0 | 0.000000071 | 0.5 | 0.27 |
| 5.26247026 | *Cc05t12380.1*<br>Transducin/WD40 repeat-like superfamily protein | 3,088 | 0.00000006 | 0.5 | 0.27 |
| **Intermediate, leaf rust incidence- Response Screening** |  |  |  |  |  |
| 2.22416916 | *Cc02t25100.1*<br>Nitrate regulatory gene 2 protein | 0 | 0.00000011 | 0.56 | 0.21 |
| 10.3678570 | *Cc10t04730.1*<br>C2H2-type domain-containing protein | 0 | 0.00000026 | 0.54 | 0.2 |
| 10.3747377 | *Cc10t04810.1*<br>WRKY domain-containing protein | 0 | 0.0000012 | 0.52 | 0.18 |
| 1.26450374<br>1.26450396 | *Cc01t08110.1*<br>Putative late blight resistance protein homolog R1B-16 | 0 | 0.0000016 | 0.51 | 0.18 |
| 1.29976260<br>1.29976261<br>1.29976262 | *Cc01t11280.1*<br>Conserved hypothetical protein | 2,741 | 0.0000013 | 0.51 | 0.18 |
| 5.28610373 | *Cc05t15840.1*<br>TPR_REGION domain-containing protein | 0 | 0.000000109983<br>1759798 | 0.55894552<br>7599274 | 0.2149040185<br>74301 |
| 7.3420157 | *Cc07t04860.1*<br>AAA domain-containing protein | 0 | 0.000002742786<br>867297 | 0.50003147<br>5501438 | 0.1719888335<br>69491 |
| 11.13698759<br>11.13698762 | *Cc11t03510.1*<br>RING-type domain-containing protein | 337,600 | 0.000003523691<br>1617069 | 0.49503097<br>4565757 | 0.1685661289<br>86321 |
| 4.27729192 | *Cc04t17060.1*<br>BHLH domain-containing protein | 85 | 0.000003208365<br>0234388 | 0.49690993<br>2910025 | 0.1698481894<br>82425 |
| 10.3876532 | *Cc10t04930.1*<br>SASA domain-containing protein | 2,641 | 0.000004767231<br>4207586 | 0.48890789<br>7990928 | 0.1644219035<br>21703 |

For the yield of green beans in the Premature population, a notable candidate gene was a putative *caffeine synthase 3* (*Cc09t06990.1*), which suggests a potential linkage between caffeine metabolism and bean characteristics that may influence overall yield. Other candidates across the traits and populations (Table 1) included involvement in diverse cellular processes (protein ubiquitination, signal transduction, and cell wall modification).

### 3.2 Bootstrap Forest analysis of SNP importance for agronomic traits

We analyzed variable importance from Bootstrap Forest models to further investigate the genetic architecture of the three agronomic traits and identify SNPs with the most significant influence on phenotype prediction. Figs. 3 and 4 present Manhattan plots showing the importance score ("Portion") for each SNP across the 11 chromosomes of *C. canephora* in the Premature and Intermediate populations, respectively. For the production of coffee beans in the Premature population (Fig. 3a), the most important SNPs were concentrated on chromosome 6, suggesting a region of significant influence on this trait.

**Fig. 3.**
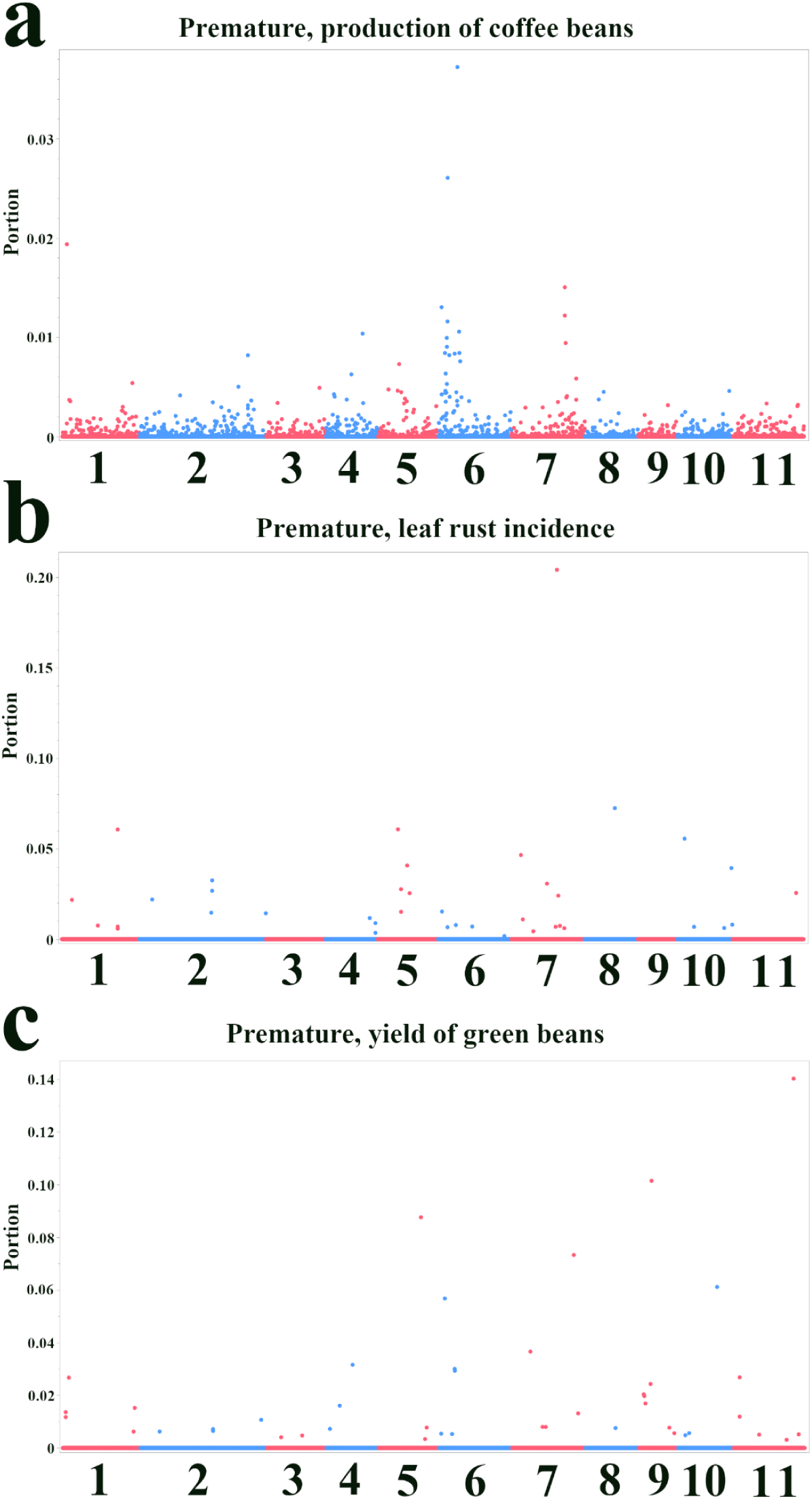
Manhattan plots showing variable importance from Bootstrap Forest models for agronomic traits in the Premature population. The x-axis indicates the chromosomal position of each SNP. The y-axis represents the variable importance score ("Portion"), which quantifies the relative contribution of each SNP to the model’s explanatory power. Higher values indicate greater importance. (a) Production of coffee beans. (b) Leaf rust incidence. (c) Yield of green beans.

**Fig. 4.**
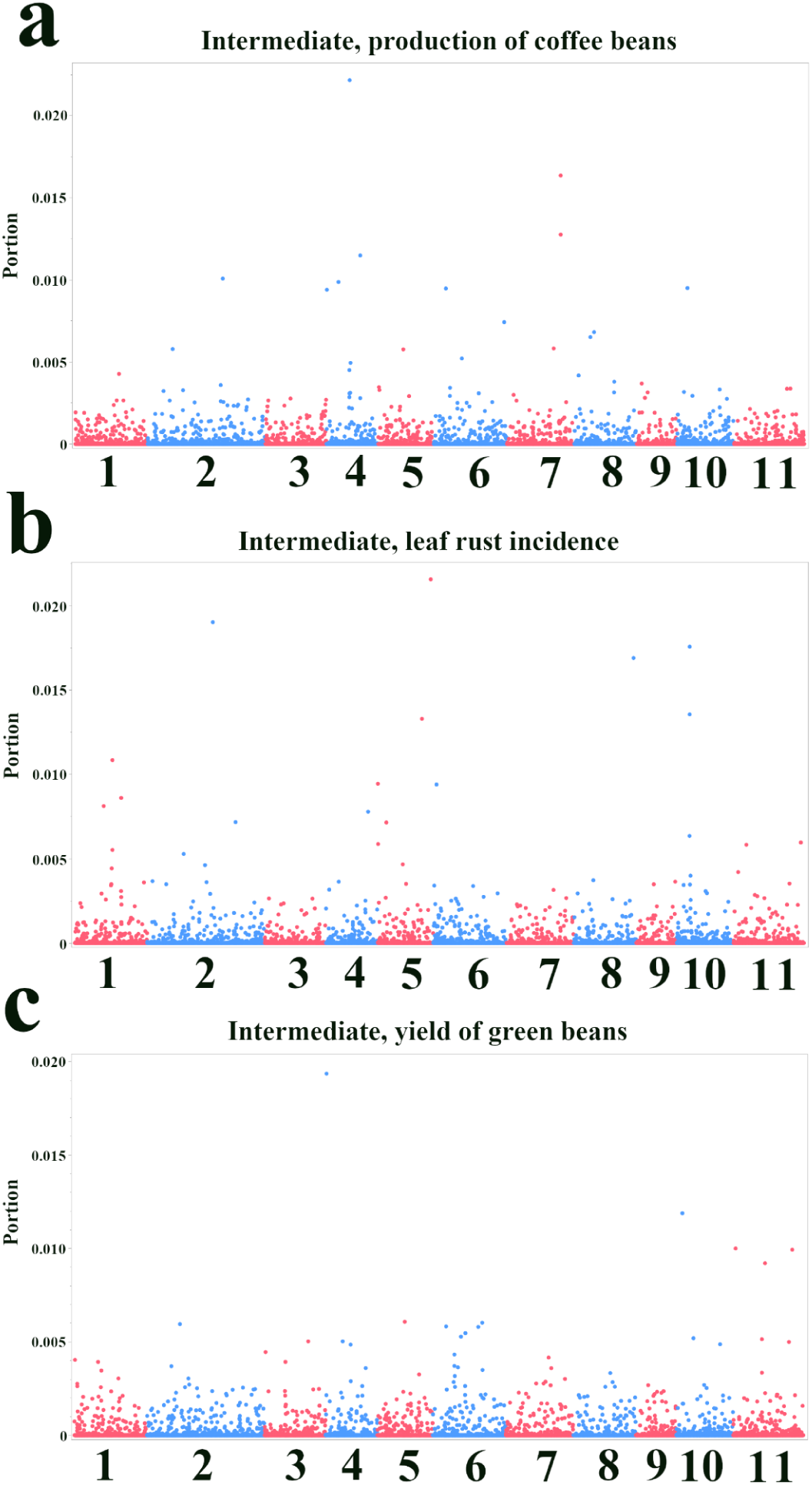
Manhattan plots showing variable importance from Bootstrap Forest models for agronomic traits in the Intermediate population. The x-axis indicates the chromosomal position of each SNP. The y-axis represents the variable importance score ("Portion"), which quantifies the relative contribution of each SNP to the model’s explanatory power. Higher values indicate greater importance. (a) Production of coffee beans. (b) Leaf rust incidence. (c) Yield of green beans.

Among the top five candidate genes in this region were those encoding an alpha/beta-hydrolases superfamily protein (*Cc06t06270.1*) and an IPPc domain-containing protein (*Cc06t03050.1*) (Table 2). For leaf rust incidence (Fig. 3b), several chromosomes exhibited SNPs with high importance scores, particularly chromosomes 5, 7, and 8. The top five candidate genes for this trait included an acyl-coenzyme A oxidase (*Cc07t15410.1*) and a hydroxyproline-rich glycoprotein family protein (*Cc08t07800.1*) (Table 2). For the yield of green beans (Fig. 3c), chromosome 11 showed a prominent SNP importance, with other important SNPs distributed across several chromosomes. A gene encoding a TORTIFOLIA1-like protein 4 (*Cc11t13960.1*) was among the top five candidates for this trait (Table 2).

**Table 2.** Top five candidate genes identified by Bootstrap Forest models for three agronomic traits in the Premature population. SNPs were ranked based on their variable importance ("Portion") in the models, with the corresponding genomic locations shown in Fig. 3. The table lists the SNP identifier (SNP ID); the nearest gene and its putative function; the distance in base pairs between the SNP and the gene’s start site (0 indicates the SNP is within the gene); and the importance score from the model.

| SNP ID | Nearest gene and function | Base pairs away | Importance score (portion) |
| --- | --- | --- | --- |
| **Premature, production of coffee beans- Bootstrap Forest** |  |  |  |
| 6.4939167 | *Cc06t06270.1*<br>Alpha/beta-Hydrolases superfamily protein | 0 | 0.037 |
| 6.2378930 | *Cc06t03050.1*<br>IPPC domain-containing protein | 172 | 0.026 |
| 1.2020393 | *Cc01t01290.1*<br>Putative 60S ribosomal protein L23a-1 | 6,605 | 0.019 |
| 7.15086633 | *Cc07t17840.1*<br>NB-ARC domain-containing protein | 19 | 0.015 |
| 6.1079450 | *Cc06t01300.1*<br>HEAT repeat-containing protein | 0 | 0.013 |
| **Premature, leaf rust incidence- Bootstrap Forest** |  |  |  |
| 7.12204503 | *Cc07t15410.1*<br>Acyl-coenzyme A oxidase | 308 | 0.2 |
| 8.21181446 | *Cc08t07800.1*<br>Hydroxyproline-rich glycoprotein family protein | 0 | 0.07 |
| 5.13342765 | *Cc05t02930.1*<br>TAF domain-containing protein | 27,160 | 0.06 |
| 1.32477500 | *Cc01t14390.1*<br>Increased DNA methylation like | 0 | 0.06 |
| 10.1733570 | *Cc10t02280.1*<br>OBG-type G domain-containing protein | 0 | 0.056 |
| **Premature, yield of green beans- Bootstrap Forest** |  |  |  |
| 11.30697733 | *Cc11t13960.1*<br>TORTIFOLIA1-like protein 4 | 0 | 0.14 |
| 9.3350785 | *Cc09t03950.1*<br>NAD(P)-binding Rossmann-fold superfamily protein | 0 | 0.1 |
| 5.24257788 | *Cc05t09820.1*<br>Glucose-1-phosphate adenylyltransferase | 1,708 | 0.088 |
| 7.20811138 | *Cc07t20130.1*<br>Protein of unknown function (DUF1365) | 26,561 | 0.073 |
| 10.20833232 | *Cc10t11950.1*<br>tRNA (guanine(37)-N1)-methyltransferase | 0 | 0.061 |

In the Intermediate population, the patterns of SNP importance differed from those observed in the Premature population (Fig. 4). For the production of coffee beans (Fig. 4a), while some SNPs on chromosomes 4 and 7 showed high importance, the overall importance scores were lower compared to the Premature population. Top candidate genes included a non-specific phospholipase C6 (*Cc04t10310.1*) and a Smr domain-containing protein (*Cc07t19900.1*) (Table 3). For leaf rust incidence (Fig. 4b), chromosomes 5 and 10 exhibited prominent peaks, with a TPR_REGION domain-containing protein (*Cc05t15840.1*) and a nitrate regulatory gene2 protein (*Cc02t25100.1*) among the top candidates (Table 3). Only a few SNPs stood out for the yield of green beans, one of which was located on chromosome 4 (*Cc04t00320.1*; Conserved hypothetical protein) (Fig. 4c; Table 3).

**Table 3.**
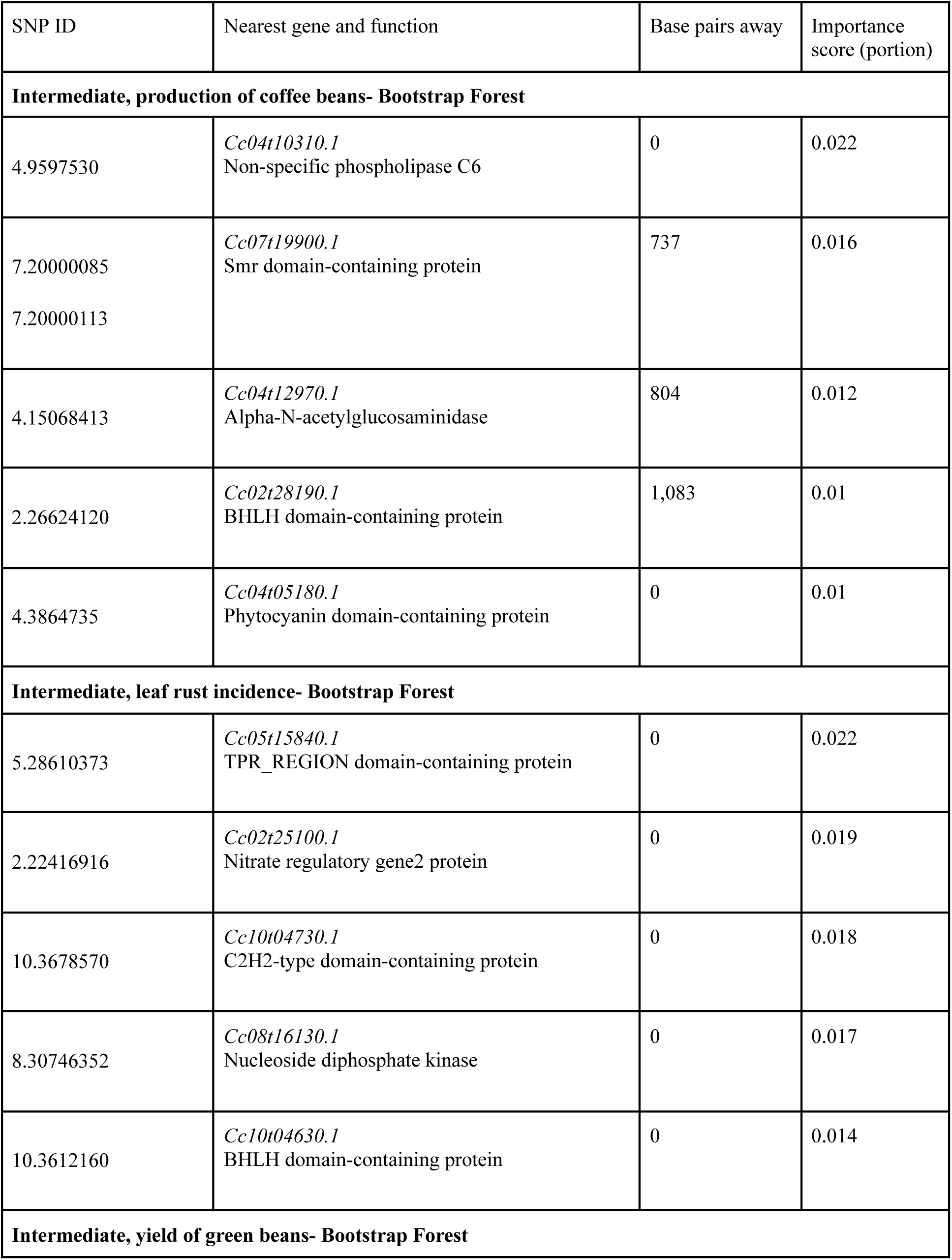
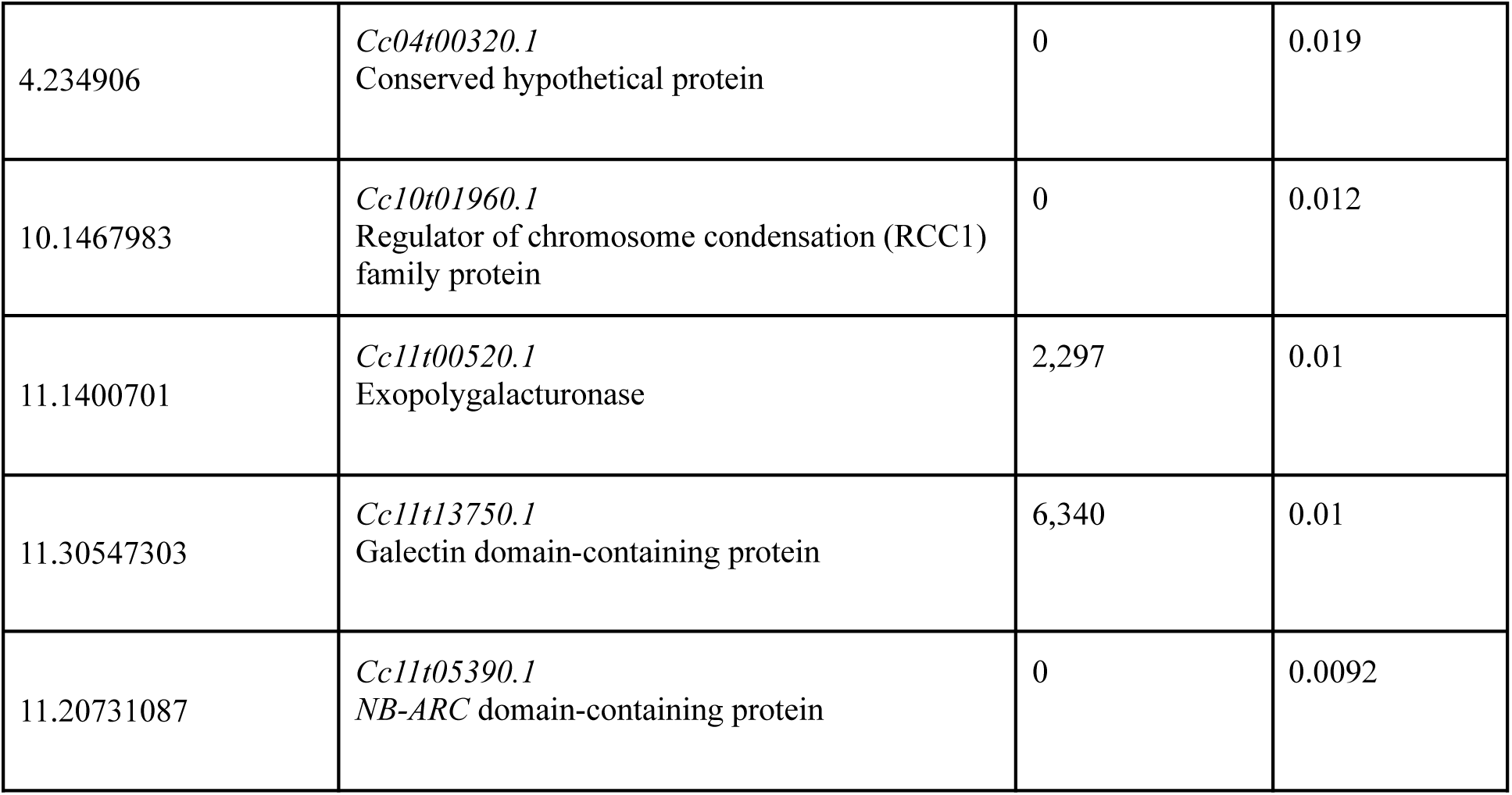
Top five candidate genes identified by Bootstrap Forest models for three agronomic traits in the Intermediate population. SNPs were ranked based on their variable importance ("Portion") in the models, with the corresponding genomic locations shown in Fig. 4. The table lists the SNP identifier (SNP ID); the nearest gene and its putative function; the distance in base pairs between the SNP and the gene’s start site (0 indicates the SNP is within the gene); and the importance score from the model.

Tables 2 and 3 list the top five candidate genes for each trait in the Premature and Intermediate populations, respectively, ranked by their importance score in the Bootstrap Forest models. It is noteworthy that some genes (*Cc06t03050.1*: IPPc domain-containing protein, *Cc05t02930.1*: TAF domain-containing protein, *Cc11t13960.1*: TORTIFOLIA1-like protein 4, *Cc02t25100.1*: Nitrate regulatory gene 2, and *Cc05t15840.1*: TPR_REGION domain-containing protein) were commonly shown as top candidates by the Bootstrap Forest models and the single-SNP analysis (see Table 1).

### 3.3 Comparative analysis and identification of consensus candidate genes

To further elucidate the biological functions underlying the distinct genetic architectures of the Premature and Intermediate populations, a GO enrichment analysis was conducted. The analysis focused on genes associated with the top 100 predictive SNPs identified by the Bootstrap Forest models for each population, assessing both individual and combined effects.

The results revealed highly distinct biological signatures for each population. In the Premature population, yield-related traits were significantly enriched for specialized cellular functions (Fig. 5a & 5b). The analysis combining all traits highlighted ’lipid modification’ (GO:0030258) and pathways related to the internal space of organelles, such as ’membrane-enclosed lumen’ (GO:0031974) and ’organelle lumen’ (GO:0043233) (Fig. 5c).

**Fig. 5.**
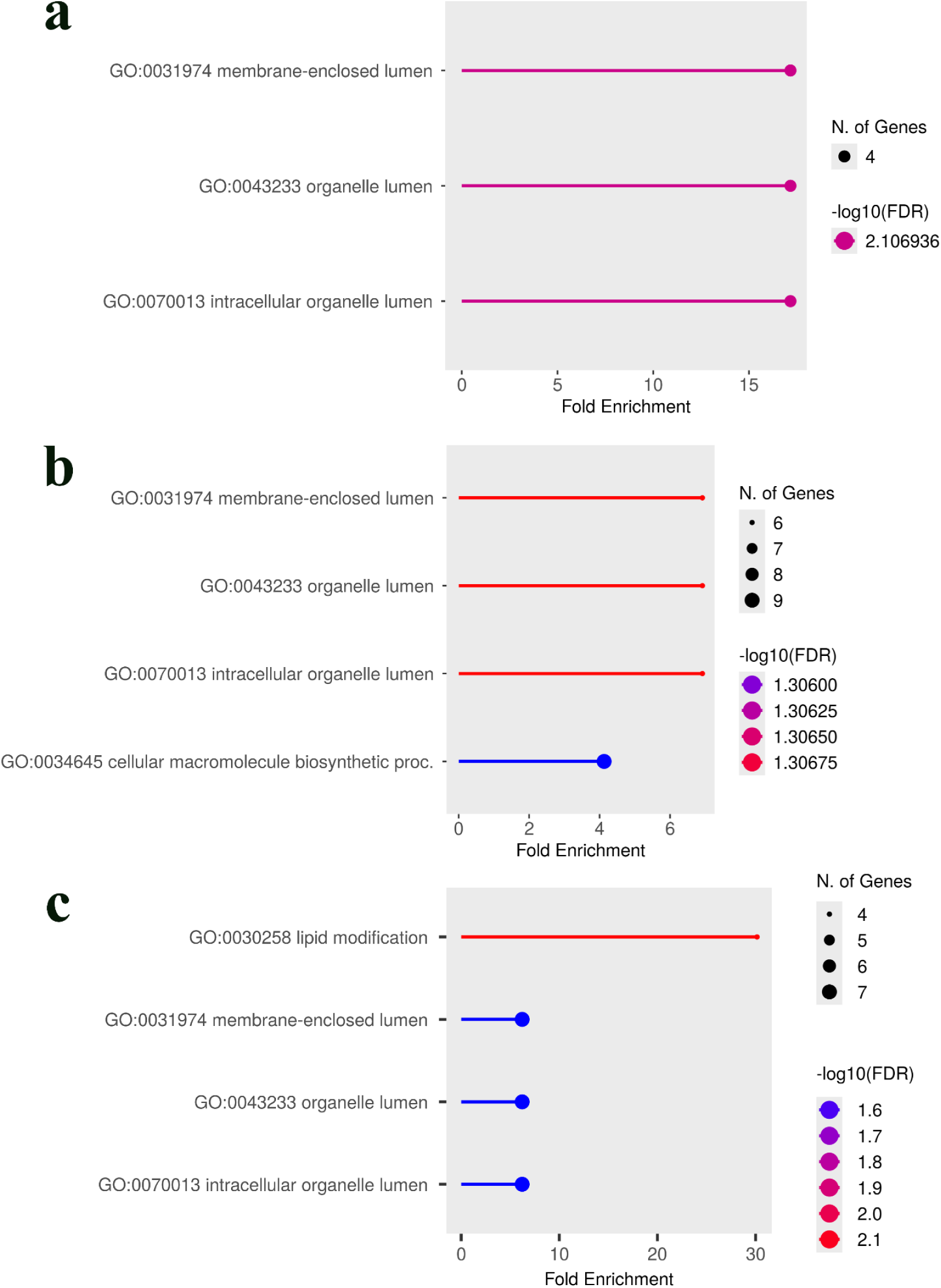
GO enrichment analysis of candidate genes in the *C. canephora* "Premature" population. The analysis was performed using genes associated with the top 100 predictive SNPs from the Bootstrap Forest models for different trait combinations. The plots illustrate the most significantly enriched biological pathways. (a) Enriched GO terms for genes associated with green bean yield, highlighting pathways related to the internal space of organelles. (b) Enriched terms for the combined traits of coffee bean production and green bean yield, which include the addition of a ’cellular macromolecule biosynthetic process’. (c) Enriched terms for the combination of all three traits (coffee bean production, green bean yield, and leaf rust incidence), identifying ’lipid modification’ as another key pathway. In all plots, the x-axis represents the fold enrichment of the term, the size of each point corresponds to the number of genes associated with the term, and the color indicates the statistical significance based on the -log10(FDR). Abbreviations used: reg., regulation; proc., process.

In contrast, the analysis of the Intermediate population showed a strong and consistent enrichment for pathways involved in regulating the actin cytoskeleton across all trait combinations. Key enriched terms included ‘reg. of actin filament depolymerization’ (GO:0030834), ’negative regulation of actin filament polymerization’ (GO:0030837), and ’actin filament capping’ (GO:0051693) (Fig. 6). Furthermore, combining yield traits introduced significant enrichment for ’response to salicylic acid’ (GO:0009751) (Fig. 6b), and the further addition of rust resistance highlighted a broader ’regulation of defense response’ (GO:0031347) pathway (Fig. 6c)

**Fig. 6.**
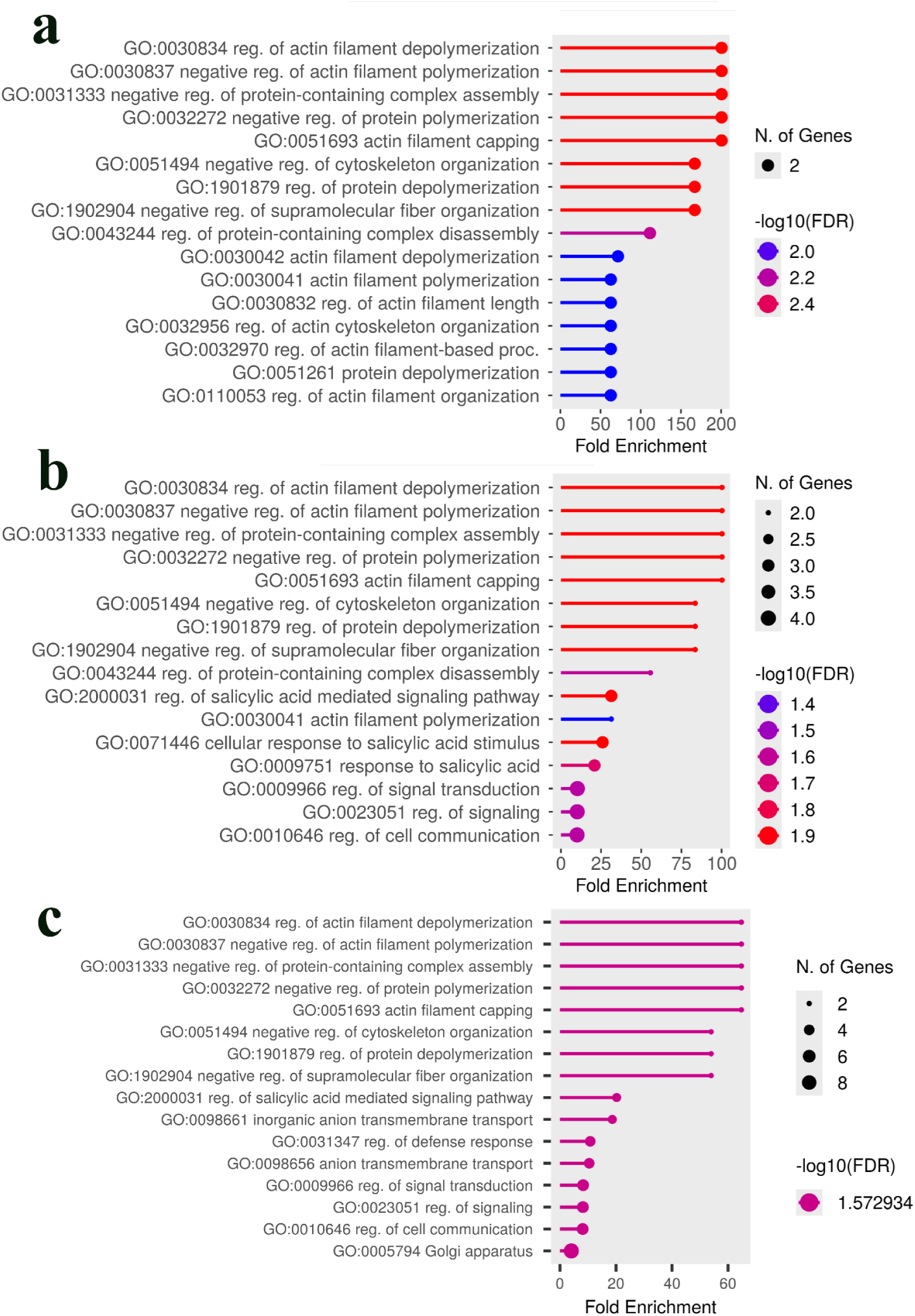
GO enrichment analysis of candidate genes in the *C. canephora* "Intermediate" population. The analysis was performed on genes associated with the top 100 predictive SNPs from the Bootstrap Forest models for various trait combinations. (a) For the green bean yield trait, the analysis reveals a strong enrichment for pathways involved in the regulation of the actin cytoskeleton. (b) Combining coffee bean production and green bean yield traits retains the actin-related terms and introduces significant enrichment for signaling pathways, including ’response to salicylic acid’. (c) The analysis of all three traits combined (coffee bean production, green bean yield, and leaf rust incidence) reinforces the importance of defense-related signaling (’salicylic acid mediated signaling pathway’, ’regulation of defense response’) alongside the persistently significant actin regulation pathways. In all plots, the x-axis represents the fold enrichment, the point size corresponds to the number of genes, and the color indicates statistical significance based on the -log10(FDR). Abbreviations used: reg., regulation; proc., process.

To identify overarching biological themes shared between the two populations, a combined GO analysis was performed by pooling the gene lists. Notably, while neither population showed significant GO term enrichment for rust resistance individually, the combined analysis revealed a strong and statistically significant enrichment for ‘chitinase activity’(GO:0004568), ‘chitin metabolic proc.’ (GO:0006030 & 0006032), and ‘programmed cell death’ (GO:0034050) (Fig. 7a). This indicates a shared, high-level defense function that becomes apparent only when data from both populations are pooled. For yield-related traits, the combined analysis was predominantly characterized by pathways involved in actin filament regulation (Fig. 7b, c), demonstrating that this strong theme from the Intermediate population defines the combined genetic signal. Interestingly, when all traits from both populations were combined, the enrichment profile shifted to highlight different pathways involved in phosphatidylinositol metabolism and sulfate transmembrane transport (Fig. 7d).

**Fig. 7.**
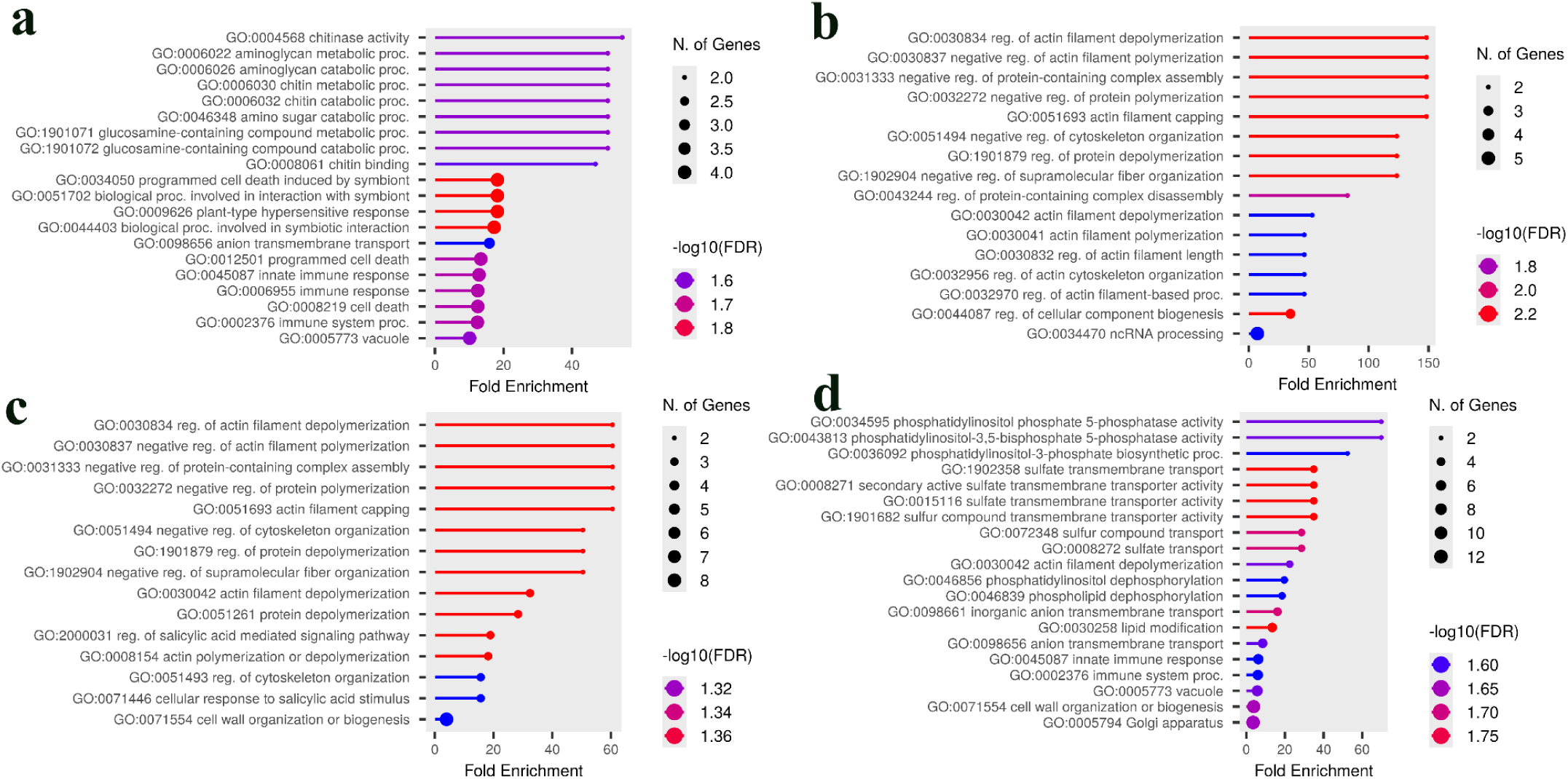
GO enrichment analysis of candidate genes from the combined "Premature" and "Intermediate" populations. The analysis was performed on pooled gene lists associated with up to the top 100 predictive SNPs from the Bootstrap Forest models in both populations. **(a)** The combined analysis for leaf rust resistance reveals a strong enrichment for defense-related pathways, including ’chitinase activity’ and ’immune response’. **(b)** For the combined green bean yield trait, the analysis is characterized by a strong enrichment for pathways related to ’actin cytoskeleton regulation’. **(c)** The addition of coffee bean production traits retains the dominant actin regulation theme while also introducing pathways related to ’salicylic acid-mediated signaling’. **(d)** When all three traits (yield, green bean, and rust) are combined, the enrichment profile highlights pathways involved in ’phosphatidylinositol metabolism’ and ’sulfate transmembrane transport’. In all plots, the x-axis represents the fold enrichment, the point size corresponds to the number of genes, and the color indicates statistical significance based on the -log10(FDR). Abbreviations used: reg., regulation; proc., process.

## 4 Discussion

Our study provides a new layer of biological interpretation that complements and expands upon previous work that established the potential for genomic prediction [16] and dissected trait stability [17] in these important *C. canephora* populations. Whereas prior analyses focused on the statistical accuracy of predictive models or the predictability of stability metrics across environments, our primary objective was to dissect the functional basis of the underlying genetic architecture. By integrating single-SNP regression, machine learning for variable importance, and a novel comparative GO framework, our work moves beyond statistical observation to mechanistic insight. The central finding of this study, that the ’Premature’ and ’Intermediate’ populations employ fundamentally different biological strategies to achieve agronomic performance, provides a new biological context for breeding efforts that was previously absent from the literature on these populations. These contrasts indicate that population-aware interpretation is essential: the same trait class can be underpinned by specialized metabolism in one breeding population and by core cellular machinery in another.

Our analysis confirms that the Premature and Intermediate populations possess distinct genomic architectures. This difference is likely attributable to their unique selection histories, which can alter allele frequencies and narrow the genetic base over time [27–29]. Additionally, potential differences in linkage disequilibrium patterns [30–32], which have been empirically shown to differ between populations based on breeding history [32, 33], and founder effects from their establishment, which are known to shape the genetic makeup of breeding populations [35], may persist and contribute to these population-specific genetic effects [36]. While our single-SNP analysis highlighted a greater number of significant associations in the Premature population, our new pathway-level analysis provides a deeper biological explanation for these differences, moving beyond statistical observation to mechanistic interpretation.

The GO analysis revealed that the two populations employ fundamentally different biological strategies to achieve their agronomic traits. In the Premature population, yield-related traits were significantly enriched for specialized cellular pathways, including ’lipid modification’ and pathways related to the ’membrane-enclosed lumen’ of organelles. This is biologically significant, as the vacuole is the primary site for sugar and metabolite storage, and its proper function is crucial for determining overall plant yield [37,38]. Indeed, direct manipulation of vacuolar transporters has been shown to increase seed yield in model plants [39]. This pathway-level finding strongly supports our identification of a putative *caffeine synthase 3* as a key candidate gene for green bean yield, as the biosynthesis and accumulation of purine alkaloids like caffeine is a key feature of seed and fruit development in *Coffea* [40–43]. In stark contrast, the genetic architecture of the Intermediate population was defined by the GO enrichment of pathways related to ’actin cytoskeleton regulation’. The actin cytoskeleton is fundamental for coordinating all aspects of plant growth, including cell expansion, intracellular transport, and the maintenance of structural integrity [44]; therefore, its enrichment suggests that genetic variation in the Intermediate population influences yield through the overall efficiency of core cellular machinery. This aligns with our identification of candidate genes such as *NPC6*, a non-specific phospholipase C known to be integral to lipid metabolism and seed oil production [45–47], and TPR_REGION domain-containing proteins, which function as core scaffolds for protein-protein interactions within essential cellular complexes involved in protein transport and hormone-mediated stress responses [48–52]. Taken together, the enrichment of organelle lumen and lipid modification pathways in the Premature population and actin-cytoskeleton regulation in the Intermediate population point to distinct routes that may converge on sink strength and carbohydrate partitioning.

A similar divergence in strategy was observed for leaf rust resistance. Our GO analysis of the Intermediate population highlighted the ’salicylic acid signaling’ pathway, linking its genetic architecture to a known hormonal defense response network that is critical for systemic acquired resistance against biotrophic pathogens [53–56]. This provides a biological context for candidate genes like the nitrate regulatory gene 2, suggesting a connection between nutrient status and defense signaling in this population [57]. In contrast, the Premature population’s resistance was associated with classical defense genes like RPP13-like protein [58–61], the NB-ARC domain, which functions as an essential signaling switch in NLR immune receptors [62–65], and CERK1, a pattern-recognition receptor that is essential for perceiving fungal chitin to trigger PAMP-triggered immunity [66–69]. The power of a comparative approach was most evident when data from both populations were pooled; only then did a shared, overarching ’defense response’ theme become statistically significant. This demonstrates that while the specific genetic components differ, they contribute to a common biological function for rust resistance across a broader genetic background. Furthermore, the final combined analysis highlighted pathways of ’phosphatidylinositol metabolism’ and ’sulfate transport’ (Fig. 7d), suggesting that fundamental membrane signaling and nutrient transport may represent a core biological system underpinning the general fitness and performance across all measured traits. Notably, a chitin/chitinase–programmed cell death module only reached significance when gene lists were pooled across populations, suggesting a convergent downstream immunity axis that single-population analyses may miss. This helps reconcile the Premature population’s CERK1/NLR emphasis with the Intermediate population’s salicylic acid–signaling enrichment.

Our results both complement and expand upon the recent polygenic GWAS analysis by Ferrão et al. [17]. While their Bayesian Sparse Linear Mixed Model (BSLMM) identified major QTL for traits like leaf blight and plant architecture, our use of complementary methods and a population-specific focus provides deeper biological context. Where our findings converge on similar genomic regions, our pathway-level analysis offers a mechanistic hypothesis for why these regions are important, a strategy that has proven effective for revealing molecular mechanisms when integrating GWAS with other functional data [70, 71]. The novel identification of distinct, pathway-level strategies (specialized metabolism vs. core cellular machinery) is a key contribution that builds upon these previous genomic studies by explaining the biological nature of the genetic variation. Our work also aligns with other studies confirming the complex, polygenic nature of traits like rust resistance in perennial crops such as *C. canephora* [72–78].

This study is not without limitations. The machine learning models were developed without an independent validation dataset due to sample size constraints, meaning the results demonstrate explanatory power rather than confirmed predictive ability. This is a critical consideration, as robust cross-validation is essential for accurately assessing and comparing the performance of genomic models [79–81]. Furthermore, we can only demonstrate statistical associations between SNPs and traits, and not definitively prove causation, a common challenge in moving from GWAS loci to causal genes [82,83]. The candidate genes we’ve identified are promising targets, but functional validation is essential to confirm their roles, a critical next step for all discovery-based genomic studies [78,83–86].

The findings from our GO analysis, however, provide clear, targeted avenues for future work. Functional validation should now prioritize not only single candidate genes like *caffeine synthase 3* in the Premature population, but also key regulators within the actin filament polymerization and salicylic acid signaling pathways now identified in the Intermediate population [84]. A deeper investigation of genotype-by-environment interactions, a well-established challenge in coffee breeding, is also crucial [9,16]. By integrating complementary analytical approaches with pathway-level analysis, this work provides a more nuanced, population-specific understanding of the genetic basis of key coffee traits and paves the way for more efficient and targeted coffee breeding programs.

## 5 Conclusion

Our comparative genomic analysis revealed significant differences in the genetic architecture of key agronomic traits between the ’Premature’ and ’Intermediate’ *C. canephora* populations (Fig. 8). These differences manifest as distinct biological strategies: the Premature population’s traits are linked to specialized metabolic pathways, including lipid modification and processes within the organelle lumen, while the Intermediate population’s traits are governed by variation in core cellular machinery, such as actin cytoskeleton regulation and salicylic acid signaling. This pathway-level understanding provides a new biological context for the specific candidate genes identified for disease resistance (e.g., *RPP13*-like, NB-ARC, *CERK1*) and caffeine biosynthesis (*caffeine synthase 3*). Further analysis with broader germplasm collections across more diverse environments is necessary to generalize these findings. Nevertheless, this study moves beyond identifying lists of candidate genes to revealing fundamentally different, population-specific biological routes to agronomic performance. Understanding these distinct strategies provides a powerful foundation for designing more targeted and efficient breeding programs in *C. canephora*.

**Fig. 8.**
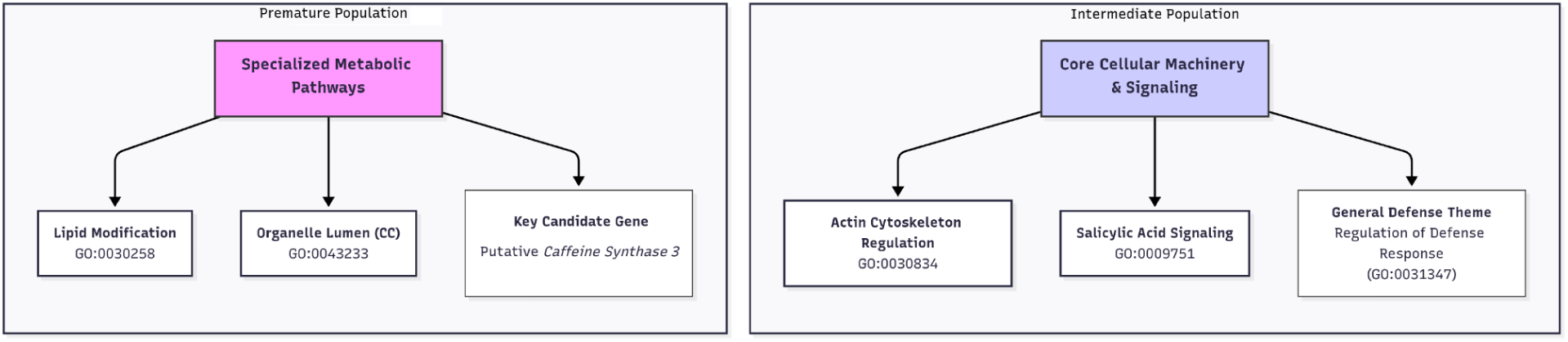
A conceptual model summarizing the distinct biological strategies for agronomic performance in the ’Premature’ and ’Intermediate’ *Coffea canephora* populations. The model visually contrasts the key findings from the comparative genomic analysis. The genetic architecture of the Premature population is linked to specialized metabolic pathways, including “lipid modification” and the cellular component "organelle lumen," and is highlighted by a key candidate gene, a putative *caffeine synthase 3*. In contrast, the Intermediate population’s traits are governed by variation in core cellular machinery & signaling, with significant enrichment for pathways like "actin cytoskeleton regulation" and "salicylic acid signaling." This figure illustrates that the two populations achieve agronomic success through fundamentally different, population-specific biological routes.

## Data availability

The raw phenotype and SNP data used in this study are available in Ferrão et al. (2019) https://doi.org/10.5061/dryad.1139fm7. The complete GO enrichment results generated and analyzed during the current study are provided as Supplementary Data 1.

## Code availability

Not applicable.

## Acknowledgements

We are also grateful to the reviewers for their constructive feedback. Mention of any trade names or commercial products in this article is solely for the purpose of providing specific information and does not imply recommendation or endorsement by the U. S. Department of Agriculture. USDA is an equal opportunity provider and employer, and all agency services are available without discrimination.

## Contributions

E.A. designed and supervised works. E.A., S.P., J.B., and S.L. analyzed experimental data. S.P., J.B., and S.L. validated the results. S.P. and J.B. visualized the data. E.A. wrote the original draft. S.P., J.B., S.L., and L.M. revised the manuscript. All authors read and approved the final manuscript.

## Ethics declarations Ethics approval

Not applicable.

## Consent to participate

Not applicable.

## Consent for publication

Not applicable.

